# An open-sourced 3D printable microscope with a large CNC stage

**DOI:** 10.1101/2024.12.31.630915

**Authors:** Clément Dégut, Michael J Plevin

## Abstract

3D printed laboratory tools are a fast-developing field, and microscopes in particular have seen several high-quality projects reported in recent years. However, currently available projects for 3D printed microscopes have not been optimised for molecular biology applications. We present an open-source, 3D-printable optical microscope that has been specifically designed for affordability, ease of use, and versatility in molecular biology applications, particularly for fluorescence detection, time lapse imaging, and higher throughput scanning of microtitre plates. The microscope integrates a large computer numerically controlled (CNC) stage, allowing precise scanning of wells in standard microtiter plates, and is equipped with a Raspberry Pi HQ camera module and customizable optical components, including standard microscope objectives. The modular design allows for a range of illumination setups, such as a 3 W blue LED for basic epifluorescence, and supports additional modifications, including dark field or RGB matrix lighting. The design opens new possibilities for cheap home-built microscopes for bioscience applications that are modular and easily adaptable for specific user needs.

## Introduction

3D printing technology has brought many high-quality affordable tools to the laboratory, from simple sample holders and magnetic racks (a large resource can be found at https://3d.nih.gov/) to much more advanced projects such as high accuracy micropipettes. In recent years, several projects have emerged that focus on affordable, high-performance, 3D-printable microscopes. Notably, the OpenFlexure project (Cicuta *et al*., 2020), which serves as the primary inspiration for the work we report here, is a fully motorized and automated microscope built around a Raspberry Pi camera. Similarly, the PUMA microscope (Tadrous, 2021) stands out for its modular design and high level of customizability. Such tools are highly needed as microscopes, despite being everyday tools in many laboratories, remain very expensive, often lack basic functionality such as the possibility to take pictures or videos, and are not easily adaptable for specific user applications.

Here, we report a design for a 3D printed optical microscope that is built around a large computer numerically controlled (CNC) stage. The design prioritizes low cost, ease of printing and assembly, and user-friendliness. The key distinction between our microscope and other designs is its ability to accommodate and fully scan standard microtiter plates or any sample up to 80 × 140 mm. It also uses the latest 12.3 MegaPixel (MP) Raspberry Pi HQ camera as sensor module. Sample illumination is provided by a simple single distant light emitting diode (LED) that can be focussed with a lightweight condenser. An additional off-axis blue LED has been included to allow fluorescence microscopy for samples labelled with compatible dyes or fluorescent proteins. Finally, an addressable RGB LED ring allows more complex illumination patterns such as darkfield and oblique illumination. We showcase several potential uses of the microscope in bioscience research, including live/dead cell assays, GFP expression tests, production of liquid-liquid phase separation (LLPS) phase diagrams, and screening of protein crystallisation conditions.

Importantly, production of the microscope does not require specialist tools or manufacturing facilities. The whole system can be assembled using only basic tools, a 3D printer and soldering iron. The cost of building of the microscope, excluding lenses, is on the order of £450 to £500 in the UK depending on which suppliers are used. The overall cost could be as low as £220 if longer delivery times can be tolerated. Full details of schematics, 3D printing and assembly instructions are available via the project GitHub page (https://github.com/cdegut/Mikroskop).

### Hardware description

The microscope design comprises two main components: (a) an optical stack that uses the Raspberry Pi HQ camera sensor with a multimodal illumination system (Figure 1); and (b) a large automated CNC stage that allows movement across a 80 × 140 mm field of view and that can accommodate standard microtitre plates (up to 35 mm thick with condenser light and 70 mm without).

**Figure 1.**
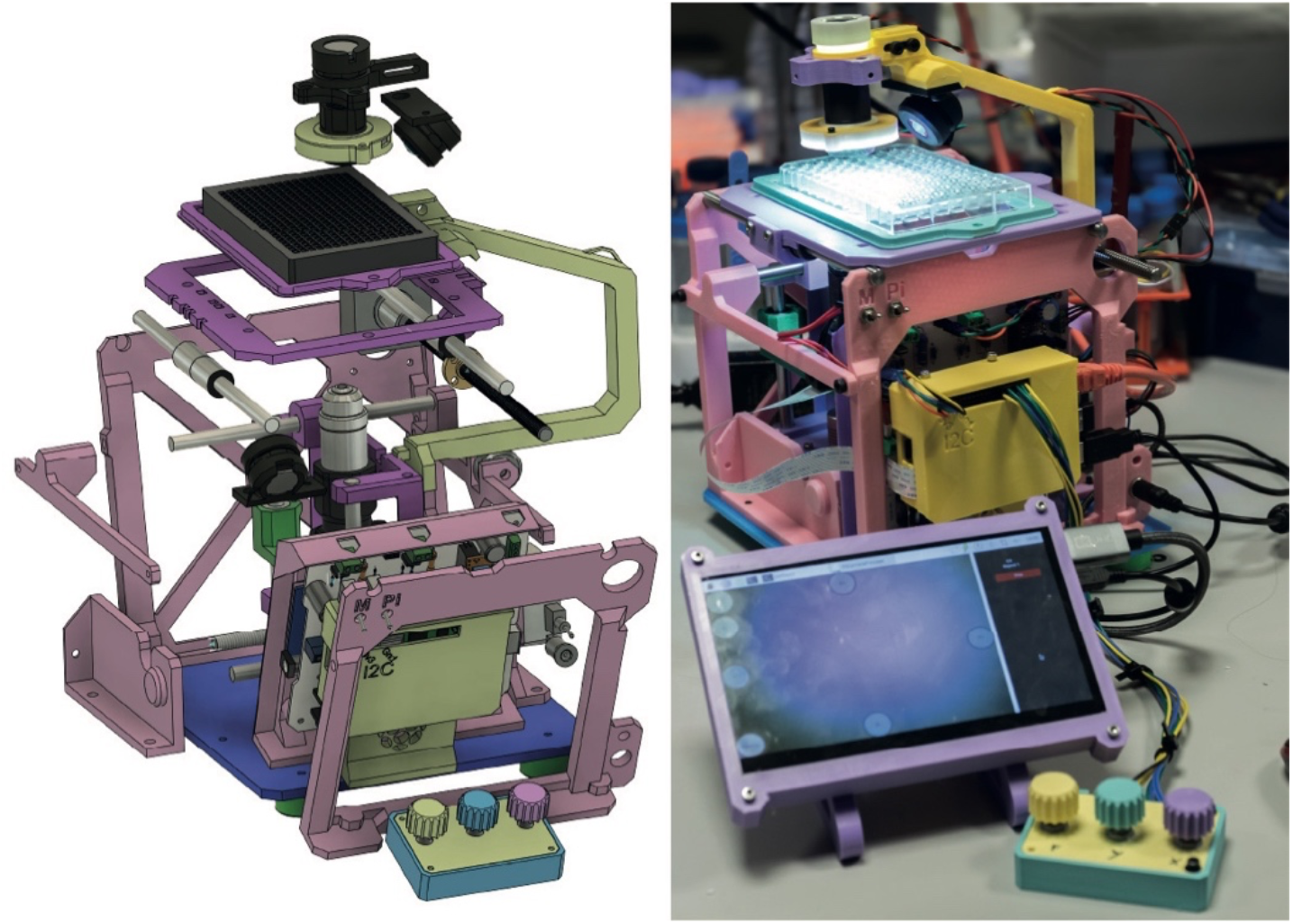
Overall microscope design. Left, schematic of design of the microscopy showing an exploded view of the overall system. Right, photo of the instrument *in situ*.

### The optical stack

The top of the optical stack houses the illumination system (Figure 2A, top). The standard configuration uses a single 3.3 V LED positioned 9 cm away from the main observation plane that shines through a 5 mm aperture. While this distant LED setup is inefficient, it offers a cost-effective way to achieve uniform sample illumination (Figure 2B top left). For higher magnification, improved contrast, and better background a f15 mm poly-methyl methacrylate (PMMA) lens can be added after the LED, and a condenser magnetically attached underneath. The combined f15 mm PMMA condenser lens can be adjusted in depth with a screw axis (Figure 2B top right). Additionally, an addressable ring of RGB LED can be attached to the condenser to allow for different modality of illumination such as dark field imaging (Figure 2B, bottom left) and pseudo differential interference contrast (DIC; also called Kristiansen Illumination; Figure 2B bottom right). The LED ring setup can also be used on the objective side to perform reflective illumination of non-transparent samples. Finally, an off-axis high-power 3 W blue LED enables basic fluorescence microscopy of samples labelled with compatible dyes, such as fluorescein or green fluorescent protein (GFP). This is made possible via a sensor with a Bayer filter, where fluorescence can be captured directly from the two green pixels with minimal cross-channel contamination. The illumination system is designed to be modular and accessible, enabling easy modifications and objective lens swapping as needed.

**Figure 2.**
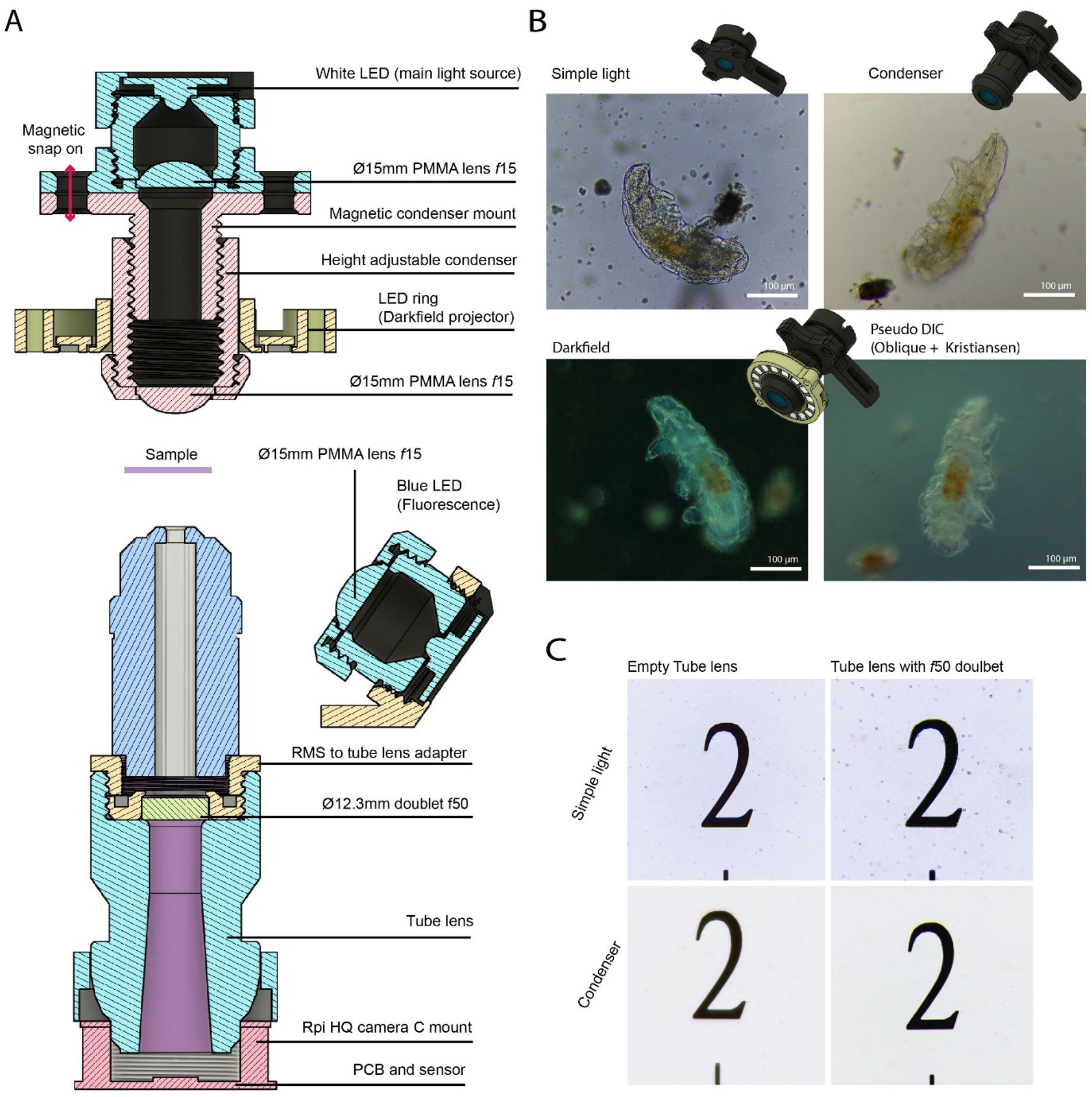
An adaptable optical set up. (A) Illumination system (top) and tube lens (bottom). The light is provided by a 1 W white LED that can be use in its own or with a magnetically attached condenser. A ring of RGB LEDs can also be added to provide darkfield illumination. The microscope uses a standard microscope lens, mounted on a Raspberry Pi HQ camera through a 3D printed tube lens. The tube lens comprises an optical doublet for field correction. (B) Example images using a 40X objective lens and different illumination modality. A tardigrade is illuminated either with just the LED (top left), the condenser (top right), the darkfield projector fully on (bottom left), or half the projector on for oblique illumination and magic tape for diffusion (i.e. Kristiansen illumination or pseudoDIC; bottom right). (C) comparison of image quality obtained using a 40X objective lens with and without the f50 doublet in the tube lens. Without condenser light (usually for low magnification work) the extra lens is not necessary. A substantial improvement in image quality is achieved when using the condenser (bottom right).

The bottom of the optical stack is a 12.3 MP RGB Bayer array sensor with a standard C-mount thread (Figure 2A, bottom). A microscope objective lens with a Royal Society of Microscopy (RMS) thread is attached to the sensor housing via a 3D-printed tube lens. An achromatic doublet lens (f = 50 mm, Ø = 12.7 mm), positioned 50 mm above the sensor, corrects the field of view (FOV) and reduces the tube length. This setup is inspired by the OpenFlexure tube lens, but the use of newer and higher resolution Raspberry Pi HQ camera offers substantial improvements in image quality. While the doublet lens significantly enhances image quality, it is not strictly necessary and can be omitted to lower costs (Figure 1C). When used in this configuration, the 12.3 MP sensor enables objective lenses to achieve their full theoretical resolution. For instance, a typical 10X objective lens with a numerical aperture (NA) of 0.25 has a theoretical maximum resolution, *d*, of 1.1 μm for green light (λ = 550 nm), as determined by Abbe’s diffraction limit formula (*d* = λ/2NA). This serves to reduce costs as a single high quality 10X lens achieves the same image resolution that a range of lenses could on a lower resolution sensor. The design of the focus axis allows the addition of filters in the infinity space of the optical system without collision.

The system is designed for an infinity corrected objective lens. To test the microscope for biosciences applications, we used long working distance (LWD) objectives. It should be noted that these are relatively expensive compared to their non LWD counterparts and that they are an extra rather than a requirement of the system. In this study we used a 10X MOTIC CCIS® Plan achromatic objective PL 10X/0.25 (WD = 16.8 mm; part number 1101001703271l; £100) and a 40X MOTIC CCIS® Plan achromatic objective LWD PL 40X/0.5 (WD = 3.0 mm; part number 1101001703261; £240).

### The CNC motion system

The second part of the system is the mechanical and motion control setup (Figure 3), which is divided into three translational axes. The X-axis moves the entire optical stack in one direction and supports the vertical F (focus) axis (Figure 3A), while the Y-axis moves the sample stage perpendicular to the X-axis (Figure 3B). The system uses off-the-shelf 8 mm linear rods and LMU-8 bearings for the rails, with movement driven by T8 trapezoidal lead screws and nuts. These are coupled to pancake-type (half-height) NEMA 17 motors connected by semi-flexible 8 to 6 mm shaft couplers. The motors complete a full rotation in 200 steps, and the drivers are configured in 8 microstepping modes, resulting in 1.6k microsteps per full rotation. Since the T-nut on the lead screw translates by 2 mm per full rotation, this provides a theoretical positional accuracy of 1.25 μm. In practice, a minimum movement size of 6.25 μm (equivalent to 5 steps) is used for the X and Y axes, which corresponds to 2 % of the field of view with a 10X lens, and single step for the focus axis. For comparison, the diameter of a single well in a 96-or 386-well plate is approximately 6 mm or 3 mm, respectively. If higher positional accuracy is required, the lead screw can be replaced with 1 mm pitch screws, and the microstepping increased to 16 though this will result in slower travel speed. In a well-constructed microscope, printed on a modern 3D printer, the axis skew (deviation from square) is less than 1 mm over the full range of motion and can be easily corrected through software adjustments (Figure 3C).

**Figure 3.**
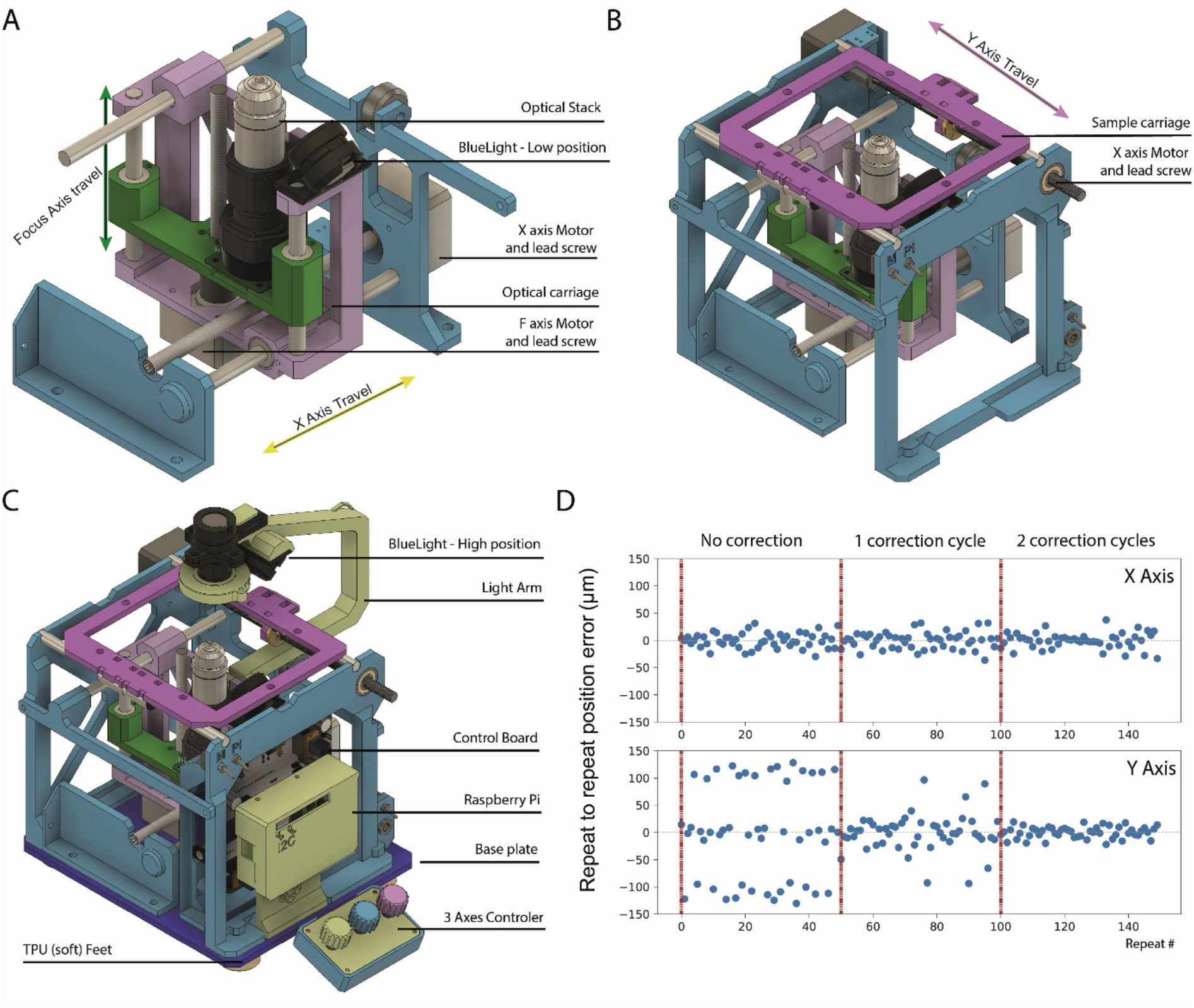
Mechanical setup of the motion system. X, Y and F axes are each composed of two linear rails and a motorised lead screw. (A) The X axis moves the optical carriage and the light arm. The focus is mounted on the optical carriage which allows the objective to move with a minimal travel distance of 1 µm. (B) The Y axis is perpendicular to X and carry the sample stage. It is mechanically independent from the other axes. (C) All components are assembled on a base plate. The microscope is supported by soft thermoplastic polyurethane (TPU) feet to absorb vibrations. (D) Three cycles of precision tuning of the X and Y axes to identify systematic errors in axis movement. The X axis (top) does not show any systematic error in reposition and the overall precision does not change following tuning. The Y axis initially shows a large error in the range ± 120 µm, but this can be rapidly reduced to a value similar to X via tuning.

The precision of the motion system (i.e. repeatability of motion) was measured as follows: the microscope was focussed on an easily recognisable feature in the FOV (in this case a crosshair) and an image was recorded. The system moved to a random position on the test axis (either X or Y) and then back to the original position at which point a second image was recorded. The FOV images were then aligned and the stage position error estimated from the image-to-image displacement. Using this method, we detected a significant systematic error on the Y axis, resulting from the stage “dragging” behind the desired motion. This error was easily corrected by splitting the dataset of measured errors into direction specific errors, taking the median of the error, and applying a direction specific motion correction equal to half of this value. The slower convergence of taking half value increases accuracy. After only two rounds of error correction the median error in Y was reduced from 176 μm (s.d. = 155 μm) to 13 μm (s.d. = 19 μm). By contrast, no similar error was seen for X-axis movement. An autocorrection script has been provided to allow users to correct for any systematic error in Y or X axis movement, and we advise that this should be performed during initial set-up of the microscope.

All non-printed components are standard off-the-shelf motion system parts that can be easily sourced from hobbyist shops or the internet. The remaining structural components are 3D-printed in poly lactic acid (PLA) at 25 % infill with little to no support material, and are designed to fit a 210 × 210 mm printing bed. PLA has been chosen due to its mechanical characteristics, its ease of printing, and because it is a bioplastic and therefore represents a more sustainable option compared to polymers derived from fossil fuels (Rezvani Ghomi *et al*., 2021). Most of the assembly uses heat-set brass inserts for plastic. The 3D-printed components are mounted on a laser-cut 5 mm acrylic plate, although this plate can also be 3D printed (we suggest 50 to 70 % triangular infill in that case). All 3D-printable parts (STL files) are available on the project GitHub page.

### Electronic main board

The electronic main board mostly serves as an interconnect for off-the-shelf modules (Figure 4). Power and control for the motors are managed by Trinamic TMC2209 stepper drivers in a StepStick-compatible format. While it is technically possible to use different drivers, the firmware employs the StallGuard4 feature of the Trinamic drivers for homing the axes. If a different driver is used, end-stop switches will need to be added, and the firmware reconfigured accordingly. The drivers are set up via UART in 8 microstepping modes with 256-step interpolation and StealthChop2 mode. The root mean square (RMS) current is defaulted to 1000 mA but can be adjusted in the firmware for compatibility with different motors.

**Figure 4.**
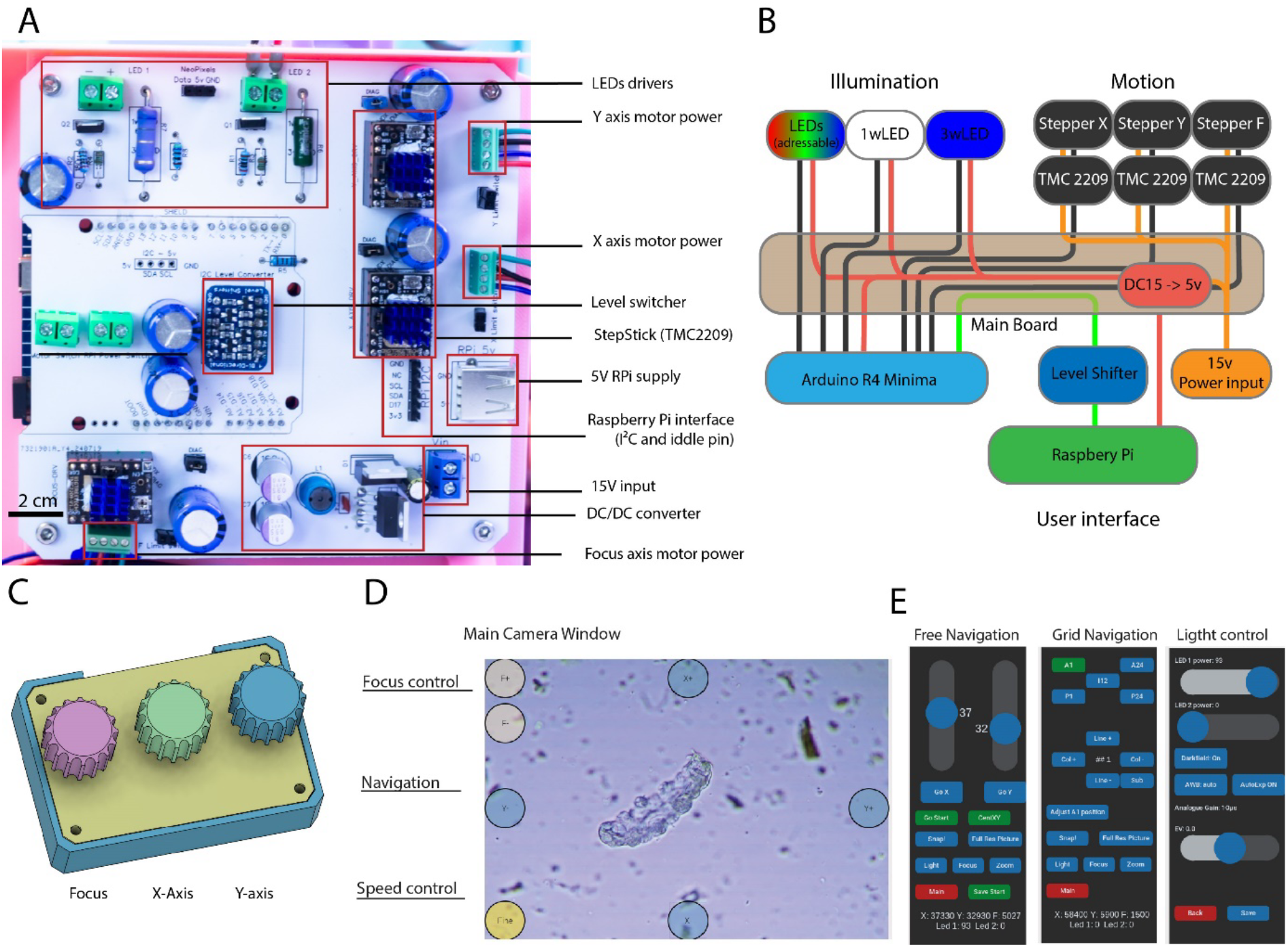
Electronic main board, and controls. (A) photograph of the populated custom PCB. (B) schematic of the interconnections. Red shows power line, black shows transistor to transistor logic (TTL), and green shows the I^2^C link. (C) The manual controller composed of three rotary encoders that provide an easy, responsive and intuitive way to control the microscope stage and focus. (D) View of the main camera window. The display shows live images while on-screen buttons allow the user to move the axes in any direction. (E) View of the side panel interface. The interface is composed of a series of side panels corresponding to different visualisation and configuration modes. The examples show “free navigation” mode, which allows free movement of either axis in any direction for large movements; “grid navigation” to move to a specific grids position that matches a microtitre plate layout; and “light control” to control both illumination and exposition.

The illumination LEDs are driven by two IRLR024NPbF gate drivers under 1 MHz pulse width modulation PWM control. This configuration reduces the bit depth resulting in a lower degree of control of the light while also giving flicker free illumination. Microstepping pulses, PWM signals, and logic are managed by an Arduino R4 Minima with custom firmware. The microcontroller receives motion instructions from a Raspberry Pi 4 via the I^2^C protocol, supplemented by a custom idle signal pin (referred to as the “ready pin” in the software). I^2^C signals pass through a 3.3 V to 5 V bidirectional level converter (BSS138). Power is supplied by a 16 V, 4.0 A power block, which is stepped down to 5 V using an LM2678T-5.0/NOPB. Components are connected on a custom double-layer PCB (1 oz/sqft copper), with all components chosen as through-hole types to eliminate the need for reflow soldering, which allows for assembly with a standard soldering iron. The firmware for the Arduino microcontroller is written in C++ and can be found in the GitHub repository under the firmware folder. It is set up as a PlatformIO project that can be uploaded to the Arduino using Microsoft VSCode. Gerber files to order the PCB are available on the project Github.

### Raspberry Pi 4 and interface

The Raspberry Pi 4 single-board computer serves as the coordinator of the system. The main software is written in Python 3, providing a hardware abstraction layer for the microscope (module “Microscope”) that defines a microscope object and enables control of the Arduino through the Raspberry Pi GPIO I^2^C pins and the idle pin (default is pin 4). The software drives the motors by sending I^2^C packets to the Arduino and checks the actual position via I^2^C requests. The camera is controlled using the Picamera2 library (https://github.com/raspberrypi/picamera2). A comprehensive graphical user interface built with PyQT and Tkinter facilitates touch input with a 7” 1024 × 600 px touchscreen that allows easy navigation within the microscope’s large 3D volume using a grid system. The interface supports image capture at both preview and full resolution, as well as video, timelapse sequences, and images on a grid. An optional physical controller can be added, consisting of three rotary encoders connected directly to the Raspberry Pi GPIO, allowing for direct control of the stage.

### Application of the system to different biological questions

Our 3D-printed microscope can image any standard or non-standard plate format. It is well suited to observe suspension or adherent cells directly in culture plates courtesy of its large stage and high-resolution sensor (Figure 5A). The 3 W blue LED enables imaging of GFP (or YFP) expression in insect or mammalian cells (Figure 5B). It is also perfectly suited for high magnification applications such as following subcellular localisation of GFP markers (Figure 5C) or observing LLPS of purified proteins, both in brightfield and fluorescence (Figure 5D).

**Figure 5.**
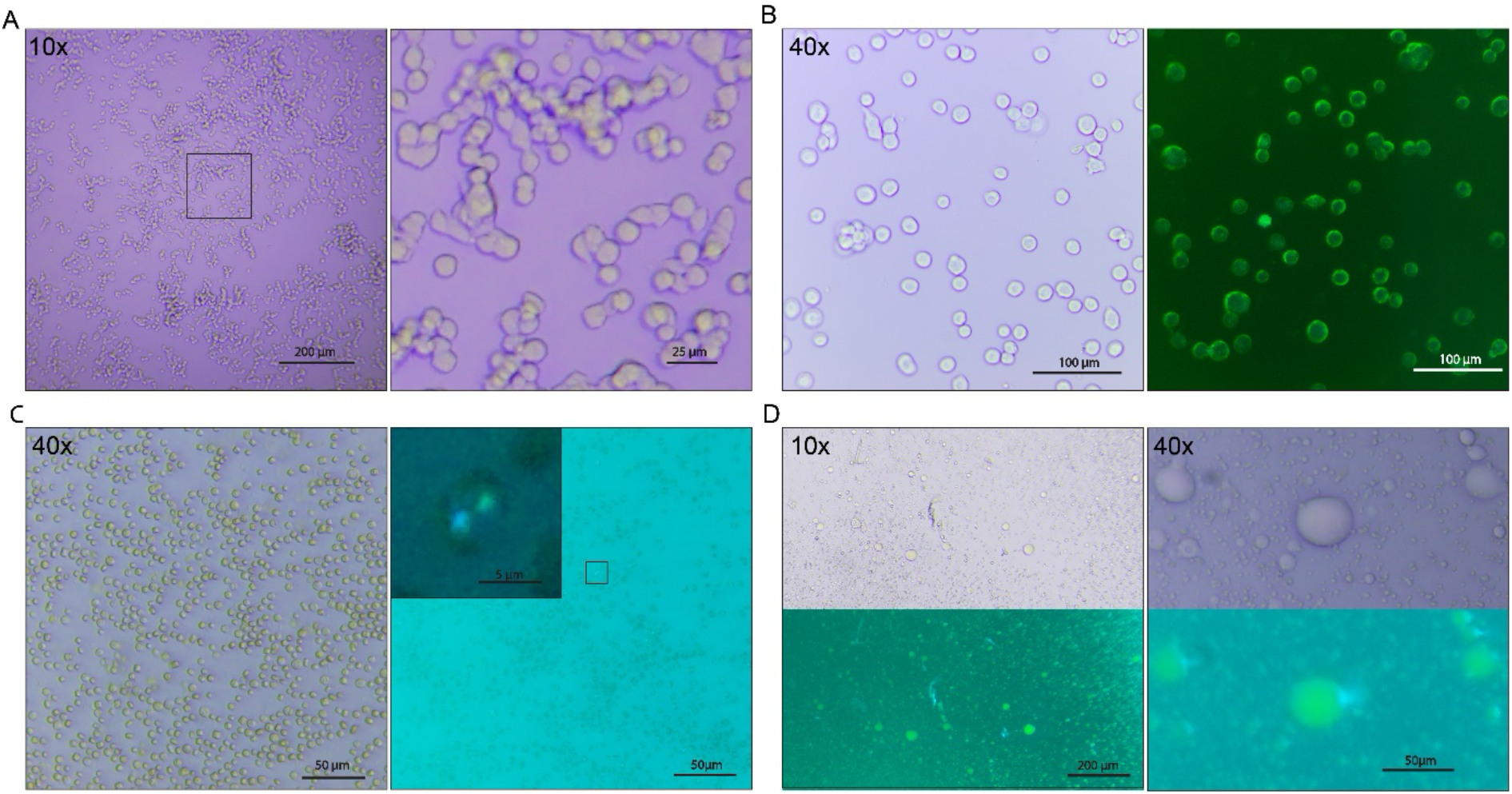
Application of the microscope in bioscience research. (A) Left, freshly split HEK293T adherent cells in D10 media in a standard 6-well plate imaged using a 10X objective. Right, the high resolution of the sensor permits a substantial digital zoom. (B) Insect cells in an 18-well culture plate expressing membrane-bound GFP imaged using a 10X objective. Left, brightfield; right, blue LED and fluorescence. The off-axis blue LED fluorescence can be used to monitor GFP expression. (C) GFP is used to monitor Rubisco localisation in diatoms. *Thalassiosira pseudonana* CCAP1085/12 cells (equivalent to CCMP1335 cells) expressing mEGFP-tagged Rubisco small subunit (Nam *et al*., 2024) were settled in an 18-well culture plate, fixed in ultra-low melting temperature agarose, and imaged using a 40X objective. Left, brightfield; right, fluorescence with zoom on a specific cell (enhanced contrast). Rubisco forms condensates in two distinct subcellular compartments (pyrenoids) that are easily distinguishable in the images collected. The high dynamic range between the two pyrenoids shows as a bleed in the blue channel for the most intense of the two spots. (D) Monitoring LLPS of SARS-CoV2 nucleocapsid protein (6 mg/mL) and a fluorescein-labelled RNA (100 μM) in 20 mM Tris pH 8.0, 50 mM NaCl. Phase separated protein-RNA droplets are clearly visible both with 10X and 40X objective lens. Preparation of SARS-CoV2 nucleocapsid protein is described in Tugaeva *et al*., 2021. All brightfield images were recorded with the condenser module on. Fluorescence images were obtained with the off-axis blue LED at full power (3 W) using only the sensor filter for detection. The objective lens used in each case is specified.

The microscope can record video up to 1,600 × 1,200 px at 30 frames per second (fps), as well as very high resolution time lapse up to 4,056 × 3,040 px at 0.5 fps (Figure 6A). The implementation of CNC motion control means that any of these modalities are available at any point on a 2D grid, and that images can be collected from multiple grid positions in sequence. A full 96-well plate can be scanned in under 5 min.

**Figure 6.**
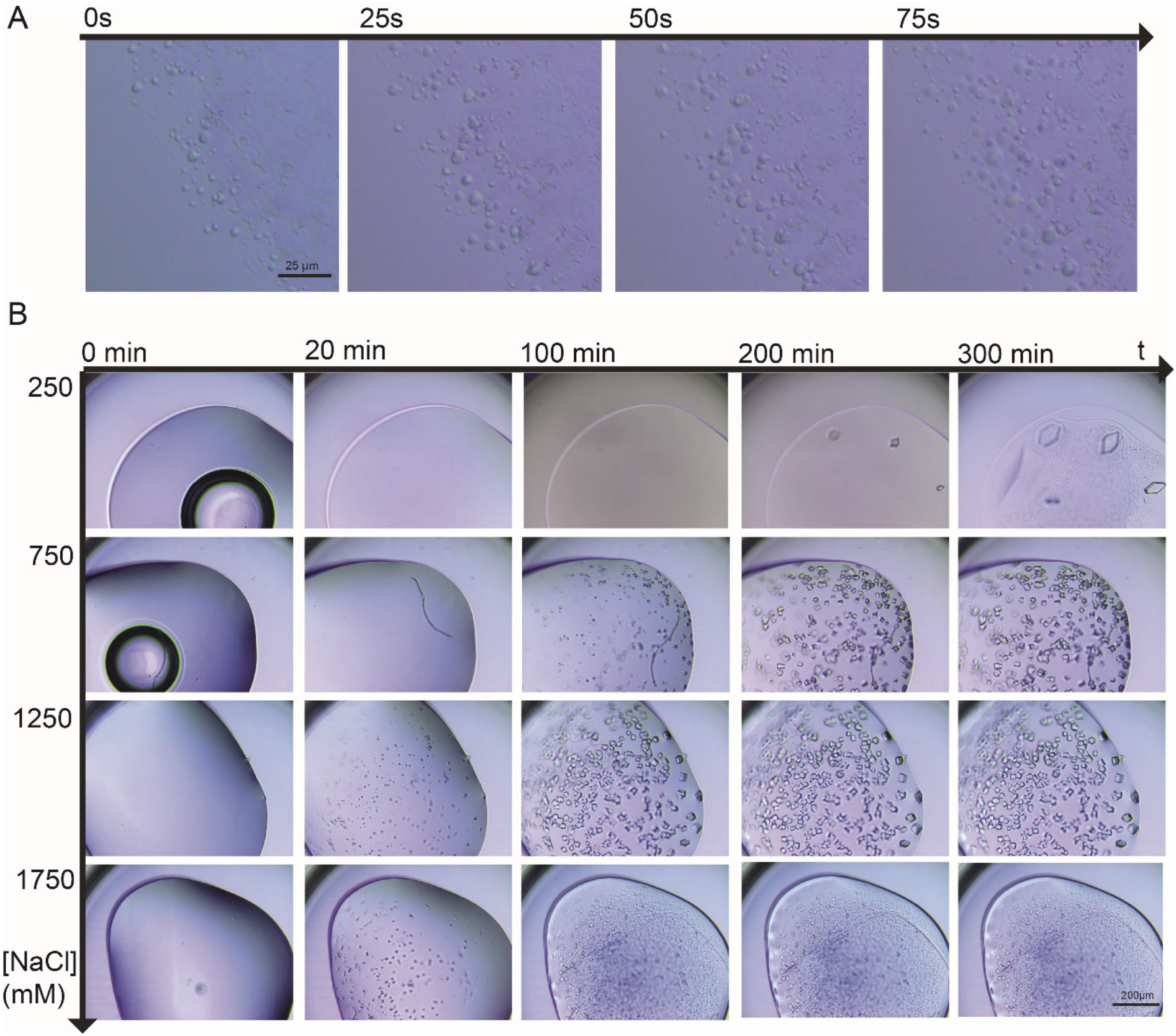
Time courses and arrays. (A) 3 µL drops of Rubisco (1 µM) and a linker protein (4 µM) in 20 mM Tris pH 8.0, 50 mM NaCl in ibidi µ-Slide 18 round wells imaged with a 40X lens. Timelapse started immediately after mixing and the sample was imaged every 5 s. Examples images from every 25 s are shown here. Protein-RNA droplets rapidly move and merge together forming larger and larger phase separated system. **B** Example of time lapse on a grid using a 10X lens. Wells in MRC 2 Lens crystallisation plate were filled with 1 µL 50 mg/mL lysozyme in 20 mM phosphate buffer, pH 4.6 and increasing concentration of NaCl (0 to 2.5 M). Wells were imaged every 5 min for 12 hours. Lower concentrations of salt lead to larger crystals while higher concentrations lead to faster crystallisation.

## Discussion

The goal of this project was to expand on recent developments in open source, low-cost 3D-printed microscopes, such as OpenFlexure or PUMA, with a new system that was designed with bioscience research applications in mind. To that end, we have designed and developed a CNC microscope using off-the-shelf electronic or 3D-printed components that has stage movement and image capture controlled by a Raspberry Pi and a purpose written user interface. Our new system has several features that set it apart from previously reported 3D-printed microscope projects. In particular, our design features a large sample stage (80 × 140 mm) that can accommodate a range of sample sizes, from microscope slides to standard-sized microtitre plates. The capacity to image microtitre plates demands a motorised stage that can automatically move to and image each well. Our design offers this capability, which represents a significant upgrade over currently available 3D-printed or low-cost microscope designs. Importantly, the inclusion of the larger stage has not compromised the quality of the optical system nor illumination capacity. We show here that our design can meaningfully image biological samples as challenging as biomolecular LLPS.

Our microscope follows open design principles, which means almost all functionality can be adapted by the user for different sample or imaging applications. For example, the sample holder design we report here is optimised for standard sized microtitre plates and slides, but it is straightforward for users to design their own bespoke magnetic attachment for any form of non-standard sample. Additionally, users can choose between many different illumination modalities including, bright field with or without condenser, off axis fluorescence, LED ring on both side of the sample, all of which are entirely compatible with the same basic microscope platform and Raspberry Pi interface. Finally, the large space on the focus axis permits addition of different modules in the infinite space of the optical path, such as filter holders and beam splitters.

It is possible to use our system to collect many different types of imaging data, ranging from standard microscopy imaging of slides to plate scanning, all while retaining the capability of high resolution microscopy. The microscope design can easily perform everyday microscopy applications in the biosciences, such as life/death cell assays, rapid imaging of microtitre plates (e.g. a 96 well plate can be fully imaged in 4-5 min), or quick assessment of cell fluorescence levels. The system can also be used for highly specific assays such as long-term time lapse of cells in culture, LLPS assays and phase diagram mapping, or screening crystallisation growth conditions. Many commercial systems offer some or all of these functions. However, the overall build cost of our system is very low (<£500, excluding lenses), making our microscope considerably cheaper than commercially available systems with the same functionality. Hence, our design is well suited to everyday applications on the lab bench as a cheap alternative to more advance microscope systems, where typically high cost or demand can limit access. One potential use of our system would be for screening and optimising experimental conditions prior to using more advanced microscopes for collecting publication quality quantitative data.

We also believe that the low cost of ease of build and modification of our microscope make it a very good educational tool across a range of STEM subjects and student ages. When fully built, the microscope can be used as a teaching tool for students learning how to use microscopes or, from parts, as a building project for students wanting to learn how to build their own scientific instruments. The open design of the microscopes means that all of the components can be easily seen by the user rather than being concealed in an opaque outer shell. As such, the design supports early training of the basic principles of microscope design and use. All of the optical elements are easy to access and modify meaning it is simple to test different imaging approaches, e.g. with or without condensers, darkfield elements, etc. The liveview and physical controller are easy to use and playful and physically connect the user to the process of image collection. The use of a Raspberry Pi for controlling the microscope and Python as the language of the interface means that students or users can write and test different operating protocols. All of the components are cheap to print or buy meaning that different adaptations can be designed and tested at reasonable cost.

We have many software and hardware developments planned for both general and specific applications. Among our current software development projects is a new multimodal imaging protocol for time lapse and grid mapping, which would allow the system to record multiple images with different settings (e.g. bright field, fluorescence, darkfield, etc) at each time point and plate position. Additionally, we are writing an on-screen scale bar overlay and a general information overlay that provides basic image properties. An autofocus procedure that would introduce the possibility of taking focus stacked images (i.e. multiple images from different focal planes) is also in development. Lastly, we are implementing an automatic tilling procedure to generate ultra-high resolution images of artificially extended FOVs.

On the hardware front, we are designing a better focus mechanism to align the condenser on the Y axis as it currently relies on shims. We have designed a complete filter holder and beam splitter assembly that would allow high quality epifluorescence; the current implementation is limited by strong illumination directionality while the bottom placement of the blue LED requires an objective lens with a working distance >3 mm. A condenser using an achromatic collector lens will also be tested to evaluate performance against the PMMA lens used in the current design. Finally, a version of the microscope where the Raspberry Pi and main board are fully separated from the main microscope platform is also under development. This modification would suit experimental applications that involve controlled environments such as cold cabinets, anaerobic chambers, or high humidity incubation chamber. Only the basic microscope would be placed under the experimental conditions with the remaining electronics and Raspberry Pi being kept at ambient conditions. Separation would help to reduce corrosion or condensation of the electronics while allowing the user to operate the microscope without having to continually perturb the controlled environment.

All plans and required documentation for building and using the microscope are available via the project GitHub page. The value of an open-source design is that our microscope is a project in constant evolution. As well as the adaptations and improvements we are planning, we hope that others will take up and use this instrument and participate in its development.

## Acknowledgements

This work was part funded by the EPSRC (York Physics of Pyrenoids Project; EP/W024063/1) and by the Wellcome Trust (ref: 204829) via the Centre for Future Health (CFH) at the University of York. We acknowledge Professor Ian Hitchcock and Dr Zoë Ingold for providing HEK cells, Dr Talha Safi for providing SF9 insect cells expressing GFP, Dr Belinda Bullard for providing samples that were key for the design and implementation of fluorescence capability, Dr Sher Keng for providing samples of SARS-CoV2 nucleoprotein, and members of the YP3 team for helpful discussions.

## References

Cicuta, P., Knapper, J., Nyakyi, P. T., Sanga, V. L., Mkindi, C., Vodenicharski, B., Dale, S., Mduda, J., Carbery, D., White, L., Lim, Z. J., Bowman, R., Mwakajinga, G. A., Baumberg, J. J., Collins, J. T., Stirling, J., Mayagaya, V., and McDermott, S. (2020). Robotic microscopy for everyone: the OpenFlexure microscope. Biomedical Optics Express, Vol. 11, 2447–2460, 11(5), 2447-2460.

Nam, O., Musiał, S., Demulder, M., McKenzie, C., Dowle, A., Dowson, M., Barrett, J., Blaza, J. N., Engel, B. D., and Mackinder, L. C. M. (2024). A protein blueprint of the diatom CO_2_-fixing organelle. Cell, 187, 5935–5950.

Rezvani Ghomi, E., Khosravi, F., Saedi Ardahaei, A., Dai, Y., Neisiany, R. E., Foroughi, F., Wu, M., Das, O., & Ramakrishna, S. (2021). The Life Cycle Assessment for Polylactic Acid (PLA) to Make It a Low-Carbon Material. Polymers, 13, 1854.

Tadrous, P. J. (2021). PUMA – An open-source 3D-printed direct vision microscope with augmented reality and spatial light modulator functions. Journal of Microscopy, 283, 259–280

Tugaeva, K. V., Hawkins, D. E. D. P., Smith, J. L. R., Bayfield, O. W., Ker, D. S., Sysoev, A. A., Klychnikov, O. I., Antson, A. A., and Sluchanko, N. N. (2021). The Mechanism of SARS-CoV-2 Nucleocapsid Protein Recognition by the Human 14-3-3 Proteins. Journal of Molecular Biology, 433, 166875

